# Efficient Gene Editing and Overexpression of Gametophyte Transformation in a Model Fern

**DOI:** 10.1101/2024.04.10.588889

**Authors:** Wei Jiang, Fenglin Deng, Mohammad Babla, Chen Chen, Dongmei Yang, Tao Tong, Yuan Qin, Guang Chen, D. Blaine Marchant, Pamela S. Soltis, Douglas E. Soltis, Fanrong Zeng, Zhong-Hua Chen

**Author notes:** Authors for Correspondence Professor Zhong-Hua Chen, ORCID ID: 0000-0002-7531-320X, Professor Fanrong Zeng.

## Abstract

The clustered regularly interspaced short palindromic repeats (CRISPR)/CRISPR-related nuclease (Cas) system allows precise and easy editing of genes in many plant species. However, this system has not yet been applied to any fern species due to the complex characteristics of fern genomes, genetics and physiology. Here, we established, for the first time, a protocol for gametophyte-based screening single-guide RNAs (sgRNAs) with high efficiency for CRISPR/Cas-mediated gene editing in a model fern species, *Ceratopteris richardii*. We utilized the *C. richardii Actin* promoter to drive sgRNA expression and enhanced CaMV 35S promoter to drive the expression of *Streptococcus pyogenes* Cas9 in this CRISPR-mediated editing system, which was employed to successfully edit a few genes (e.g., *nucleotidase/phosphatase 1, CrSAL1*; *Cryptochrome 4, CRY4*) and *CrPDS*, encoding a phytoene desaturase protein that resulted in an albino phenotype in *C. richardii*. Knockout of *CrSAL1* resulted in significantly reduced stomatal conductance (*g_s_*), leaf transpiration rate (*E*), stomatal/pore length, and abscisic acid (ABA)-induced reactive oxygen species (ROS) accumulation in guard cells. Moreover, *CrSAL1* overexpressing plants showed significantly increased net photosynthetic rate (*A*), *g_s_, E* and intrinsic water use efficiency (*iWUE*) as well as most of the stomatal traits and ROS production in guard cells compared to those in the wild-type (WT) plants. Taken together, the optimized CRISPR/Cas9 system provides a useful tool for functional genomics in a model fern species, allowing the exploration of fern gene functions for evolutionary biology, herbal medicine discovery and agricultural applications.

## Introduction

First appearing in the fossil record around 360 million years ago (MYA), true ferns form the second largest vascular plant lineage after angiosperms with more than 10,500 species (https://www.worldfloraonline.org/). These numerous species have been instrumental in shaping plant biodiversity and many ecosystems on Earth, resulting in a breadth of adaptations and innovations that are fascinating for research in genomics, evolution, ecology, molecular biology, and physiology (Cai et al., 2021; Marchant et al., 2022). Compared to other vascular plants, distinct genes (e.g., *phenolic acid decarboxylases, aerolysin-like*, and *12-oxophytodienoic acid*) might protect ferns from biotic (Pennisi, 2023; Wei et al., 2023) and abiotic stresses (Yan et al., 2019). Many fern species are used in traditional medicine for treating fevers, relaxing muscles, and relieving pain due to the active chemical compounds they produce (Cao et al., 2017; Kumar et al., 2023; Pohthmi and Sharma, 2023).

CRISPR/Cas has been widely used in plant molecular research due to its simplicity, versatility, and efficiency for gene editing (Xie et al., 2015; Endo et al., 2019; Wang et al., 2020; Cardi et al., 2023). The cellular repair of CRISPR/Cas-mediated double-strand breaks by non-homologous end joining using sgRNA and Cas nuclease can lead to the modification of genes (Wang et al., 2018; Wang et al., 2020). The ability to reprogram CRISPR/Cas with engineered sgRNA to target any gene of interest allows plant scientists to develop new plant varieties with desired traits and reducing the regulatory complication of genetically modified organism (GMO) (He et al., 2022; Cardi et al., 2023; Pacesa et al., 2024). For instance, CRISPR/Cas-mediated inactivation significantly enhanced grain weight in rice (*Oryza sativa*) by targeting *OsGW5* (Liu et al., 2017) and *OsMADS1* (*MADS-BOX TRANSCRIPTION FACTOR 1*) (Wang et al., 2024), production of low-gluten wheat (*Triticum aestivum*) through editing the *α-gliadin* gene array (Sánchez-León et al., 2018), and powdery mildew resistance of tomato (*Solanum lycopersicum*) (Nekrasov et al., 2017). In the past decade, CRISPR/Cas technology has been successfully utilized to modify more than 130 green plant species based on a recent review (Cardi et al., 2023), including 110 angiosperms (mostly agricultural and horticultural crops with significant economic values) (Kis et al., 2019; Wang et al., 2023), and 7 gymnosperms (Ren et al., 2021; Ye et al., 2023), 3 mosses (Tansley et al., 2023; Tavernier et al., 2023; Yuan et al., 2023), and 12 algae (Belshaw et al., 2023; Patel et al., 2023; Zhang et al., 2023) without any species of ferns or lycophytes.

*Ceratopteris richardii* is a fast-growing, small, tropical homosporous fern that has been used for decades as the model fern species (Marchant et al., 2019). Genetic transformation has been performed in *C. richardii* for functional genomics (Plackett et al., 2014; Plackett et al., 2015) such as discovering the roles of genes in sex determination (Youngstrom et al., 2019), genome structure, developmental biology (Plackett et al., 2018; Geng et al., 2022), hybridization and reproductive barriers (Youngstrom et al., 2022; Withers et al., 2023), and apogamy (Bui et al., 2017). In addition, the molecular function of some *C. richardii* genes have been studied through RNA interference (RNAi) (Plackett et al., 2018; Withers et al., 2023) and overexpression methods (Youngstrom et al., 2022). While the genetic transformation of fern gametophytes as the explant usually has a low success rate, it should be noted that the majority of these methods were developed and optimized according to the well-established protocols targeting to angiosperm flowers, immature embryos, and calli (Bui et al., 2015; Bui et al., 2017). Efficient gene editing protocol for fern species has not been developed, but an efficient and fast verification system in *C. richardii* will facilitate the analysis of gene function in ferns (Frangedakis et al., 2023).

Nucleotidase/phosphatase SAL1, also known as FIERY1 (FRY1) (Ishiga et al., 2017), has dual enzymatic activity of nucleotidase and inositol phosphatase, which functions largely in responses to abiotic stresses through inositol signaling and nucleotide metabolism (Jia et al., 2019). Transient silencing of *SAL1* and loss-of-function mutants led to enhanced drought tolerance in *T. aestivum* (Manmathan et al., 2013; Abdallah et al., 2022) and *Arabidopsis thaliana* (Wilson et al., 2009; Estavillo et al., 2011), while *OsSAL1* overexpression plants were severely impaired in drought tolerance of rice (Liu et al., 2023). Additionally, *GhSAL1* improved cold tolerance via inositol 1,4,5-triphosphate-Ca^2+^ signaling pathway in cotton (*Gossypium hirsutum*) (Shen et al., 2023). Our previous study showed that *C. richardii* SAL1 (CrSAL1) and its byproduct 3’-phosphoadenosine-5’-phosphate (PAP) function as chloroplast stress signals and participated in the abscisic acid (ABA) signaling pathway for drought response and stomatal regulation (Zhao et al., 2019), but *CrSAL1* was not functionally verified through genetic engineering in *C. richardii*.

Here, we established an efficient gene-editing platform for *C. richardii* transformation using gametophytes. We improved targeting and editing efficiency of sgRNAs for an optimized *Agrobacterium*-mediated CRISPR/Cas9 system via the successful editing of *CrSAL1* (*Ceric.25G052000.1*), *CrPDS* (*Ceric.08G066500.1*), *CrCRY4* (*Ceric.03G029200.1*), and *CrYSL* (*Ceric.20G086500.1*) in *C. richardii*. Knockout and overexpression of *CrSAL1* resulted in distinctive phenotypes in gas exchange parameters and stomatal traits in the transgenic plants compared to those in the WT. Our study suggests that the CRISPR/Cas system and the potentially expanded toolkit for gene editing in ferns will facilitate more rapid gene discovery and functional validation for evolutionary biology, herbal medicine, and agricultural applications.

## Results

### Selection of fern species and developmental stages for transformation

Several reference genome of ferns have been assembled in recent years, including *Azolla filiculoides* (0.75 Gb, *n* = 22), *Salvinia cucullata* (0.26 Gb, *n* = 9) (Li et al., 2018), *Alsophila spinulosa* (6.27 Gb, *n* = 69) (Huang et al., 2022), *Adiantum capillus-veneris* (4.83 Gb, *n* = 30) (Fang et al., 2022), *Ceratopteris richardii* (7.46 Gb, *n* = 39) (Marchant et al., 2022), and *Marsilea vestita* (1.0 Gb, *n* = 20) (Rahmatpour et al., 2023) (Table 1). These high-quality genome sequences enable future research into the functional genomics and applications of ferns (Chen, 2022; Kinosian and Wolf, 2022; Frangedakis et al., 2023). In the available transformation methods, particle bombardment and *Agrobacterium*-mediated stable transformation have been successfully applied to C. *richardii* (Plackett et al., 2014; Bui et al., 2015) and *Pteris vittata* (Muthukumar et al., 2013). These robust transformation methods have paved the way for developing of gene editing in ferns. While *Pteris vittata* lacks the necessary genomic information for extensive genetic manipulation (Petlewski and Li, 2019), the recent publication of the *C. richardii* genome led us to select *C*. *richardii* as the most suitable fern species for establishing a gene editing protocol.

**Table 1.** Overview of overexpression, RNAi, CRISPR/Cas9 in fern species.

| Species | Genome size | Chromosome<br>number | Transformation methods |  |  |  |  |
| --- | --- | --- | --- | --- | --- | --- | --- |
|  |  |  | RNAi | DNAi | Agrobacterium-mediated | Particle<br>bombartment | CRISPR/Cas9 |
| *Ceratopteris richardii* | 7.46 Gb, Marchant et al. 2022 | n=39 | Stout et al. 2003 | Rutherford et al. 2004 | Muthukumar et al. 2013; Bui et al. 2015 | Plackett et al. 2014, 2015 | This study |
| *Azolla filiculoides* | 0.75 Gb, Li et al. 2018 | n=22 | - | - | - | - | - |
| *Salvinia cucullata* | 0.26 Gb, Li et al. 2018 | n=9 | - | - | - | - | - |
| *Marsilea vestita* | 1.0 Gb, Rahmatpour et al. 2023 | n=20 | Klinkand Wolniak 2000 | - | - | - | - |
| *Adiantum capillus-veneris* | 4.83 Gb, Fang et al. 2022 | n=30 | - | Kawai-Tooyoka et al. 2004 | - | Kawai et al. 2003 | - |
| *Alsophila spinulosa* | 6.27 Gb, Huang et al. 2022 | n=69 | - | - | - | - | - |
| *Pteris vittata* | - | - | - | - | Muthukumar et al. 2013 | - | - |

Unlike seed plants, homosporous ferns, including *C. richardii,* possess morphologically and developmentally distinct free-living haploid gametophytes and diploid sporophytes (Figure 1A). The germination of a haploid spore to produce a photosynthetic thallus initiates the gametophytic generation. Hormonal sex determination of C. *richardii* differentiates individual gametophytes into distinct male or hermaphrodite sexes (Conway and Di Stilio, 2020). Archegonia (female gametangia) and antheridia (male gametangia) develop to produce motile sperm and eggs, respectively (Figure 1A). Only one archegonium is fertilized, resulting in a single diploid zygote per gametophyte. This first step in the diploid sporophyte generation is crucial for genetic transformation (Muthukumar et al., 2013; Bui et al., 2015; Bui et al., 2017). Extrapolating from the successful transformation of the liverwort *Marchantia polymorpha* (Ishizaki et al., 2008) and *C. richardii* (Bui et al., 2015) gametophytes via *Agrobacteria*, we developed an *Agrobacterium*-mediated gametophyte system for gene knockout in *C. richardii*. The life cycle of *C. richardii* is completed with the production of haploid spores (Figure 1A).

**Figure 1.**
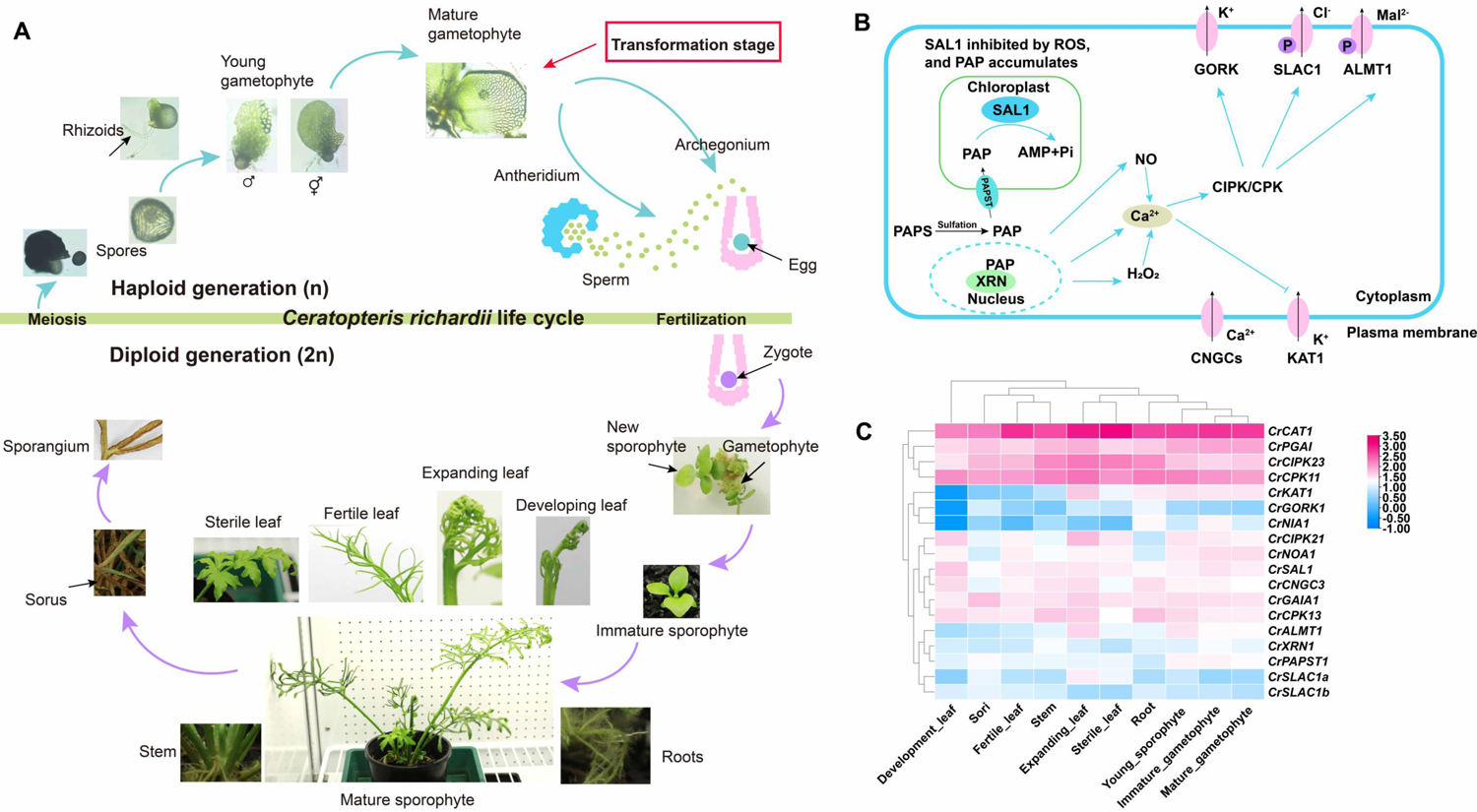
The lifecycle, proposed model and gene expression of SAL1-PAP retrograde signaling in a model fern species *Ceratopteris richardii*. The lifecycle of *C. richardii* (A). Images are not to scale. After meiosis produces haploid spores, the haploid gametophyte (n) starts generation. Spores germinate into either male gametophytes or hermaphrodite that produce gametes (sperm and egg) through mitosis. After fertilization, the diploid sporophyte (2n) generation begins as a zygote that generates into an embryo with its first root and leaf, initially dependent on the gametophyte. In the vegetative stage, the independent sporophyte produces sterile leaves (trophophyll), followed by increasingly dissected and lobed fronds. In the reproductive stage, the fertile leaves (sporophyll) of the sporophyte develop sporangia through meiosis on their undersides, closing the cycle. Model of SAL1-PAP retrograde signaling in plant (B). Expression of key genes associated with SAL1 pathway in diverse tissues such as immature gametophyte, mature gametophyte, young sporophyte, expanding leaf, development leaf, fertile leaf, sterile leaf, stem, root, sori. SAL1, 3’(2’),5’-bisphosphate nucleotidase 1; PAP, 3’-phosphoadenosine 5’-phosphate; XRN, exoribonuclease; GORK, guard cell outward rectifying K^+^ channel; SLAC1, S-type anion channel 1; ALMT1, aluminum-activated malate transporter 1; CIPK, CBL-interacting serine/threonine-protein kinase; CPK, calcium-dependent protein kinase; CNGC, cyclic nucleotide-gated ion channel; GAIA, GAMETOPHYTES ABA INSENSITIVE ON A_CE_1; CAT, catalase peroxidase; PAPST, sulfate donor 3’-phosphoadenosine 5’-phosphosulfate transporter; NOA, oxide-associated1; NIA, nitrate reductase; XRN, exoribonuclease.

### Identification and cloning of U6 promoter and Actin promoter from *C. richardii*

The core competent for CRISPR/Cas9 system contains the expression cassettes of sgRNA and the SpCas9 nuclease. Guide RNAs for genome editing have been produced using a range of Pol III promoters (Xie et al., 2015; Kor et al., 2022). We found seven *U6 small nuclear ribonucleoprotein* genes (*Ceric.17G074700, Ceric.33G040100, Ceric.09G088700, Ceric.02G026900, Ceric.1Z290000, Ceric.03G070800, Ceric.03G071600*) in the *C. richardii* genome (https://phytozome-next.jgi.doe.gov/info/Crichardii_v2_1), which showed high expression in gametophyte, leaf, stem, and root (Supplemental Figure S1A). However, the promoters of these *C. richardii* genes do not contain the upstream sequence element (USE) and TATA elements, which are the typical structural properties of the Pol III promoters (Kor et al., 2022). Therefore, we used the sequences of the *A. thaliana* U6-26 snRNA (X52528, AT3G13857) and the *T. aestivum* U6 gene (X52528, ENSRNA050022746-T1) (Poovaiah et al., 2021) sequences to compare with the upstream U6 promoter regions in *C. richardii*. We identified three promoters including CrU6-1 (Ceric.13G012200), CrU6-2 (Ceric.13G012300), and CrU6-3 (Ceric.1Z176900), which possess the USE and TATA elements (Supplemental Figure S1B). However, these genes were not highly expressed in root, stem, leaf, or gametophyte of *C. richardii* (Supplemental Figure S1A).

Previous studies showed that a single Pol II promoter (either constitutive or inducible) can also achieve effective gene editing (Hassan et al., 2021; Cardi et al., 2023) in *O. sativa* (Tang et al., 2016; Ren et al., 2019), *T. aestivum* (Luo et al., 2021), *Hordeum vulgare, S. lycopersicum, Medicago truncatula* (Čermák et al., 2017), and the diatom *Phaeodactylum tricornutum* (Taparia et al., 2022). The *Actin* promoter isolated from *P. vittata* was able to function efficiently in both *P. vittate* and *Ceratopteris thalictroides* (Muthukumar et al., 2013). A 916 bp fragment, located at the upstream of the *CrActin* was isolated and considered as the putative promoter (Supplemental Figure S2A), which was instead of the OsU3 promoter in pRGEB32 (Xie et al., 2015) to drive the expression cassettes of sgRNA (Luo et al., 2021). The Cas9 protein also reported to be driven by the enhanced CaMV 35S promoter (Li et al., 2013; Awasthi et al., 2021; Cui et al., 2021). Therefore, the native maize ubiquitin promoter (ZmUbi) promoter in the original construct pRGEB32 was replaced by the enhanced 35S promoter (Supplemental Figure S2B), which was designated as pRGEB32-CrActin.

### An efficient Agrobacterium-mediated transformation of *C. richardii* using hygromycin selection

To get positive transformants with gene editing or overexpression, the transformation protocol of *C. richardii* was optimized through adjusting the time for enzyme treatment, co-incubation and the concentrations with hygromycin for positive selection (Table 2). Subsequently, *CrSAL1* was selected to establish the *Agrobacterium*-mediated transformation of *C. richardii*. SAL1-PAP retrograde signaling is involved stomatal opening and closure through ROS, Ca^2+^, and nitric oxide (NO) pathways and ion channel (Pornsiriwong et al., 2017; Zhao et al., 2019) (Figure 1B). Here, we found that key component of the SAL1-PAP retrograde signaling pathway such as *CrSAL1, CrCAT1*, ion channels (*CrKAT1, CrALMT1, CrCNGC*) and protein kinases (*CrCIPK11, CrCIPK23*) displayed high levels of expression in most of the tissues, particularly leaves (Figure 1C).

**Table 2.** Factors affecting the efficiency of genetic transformation of the fern species *C. richardii*.

| Construct | Total number of gametophytes | OD value | Enzyme treatment time | Co-incubated *Agrobacterium* time | Hygromycin selection | Transformants | Transformation efficiency | Tested T0 seedling | Mutated T0 seedling number | Ratio |
| --- | --- | --- | --- | --- | --- | --- | --- | --- | --- | --- |
| Untransformed | 215 | 0.4 | 15 min | 15 min | 10 mg/L | 0 | 0% | - | - | - |
| *SAL1-OE* | 893 | 0.4 | 15 min | 15 min | 10 mg/L | 8 | 0.90% | - | - | - |
| *SAL1-OE* | 537 | 1.2 | 15 min | 15 min | 10 mg/L | 6 | 1.12% | - | - | - |
| *SAL1-OE* | 768 | 0.8 | 2 h | 1 h | 5, then 20 mg/L | 73 | 9.51% | - | - | - |
| *CRY4-OE* | 194 | 0.8 | 2 h | 1 h | 5, then 20 mg/L | 23 | 11.68% | - | - | - |
| *YSL-OE* | 351 | 0.8 | 2 h | 1 h | 5, then 20 mg/L | 26 | 7.41% | - | - | - |
| *GRF-OE* | 159 | 0.8 | 2 h | 1 h | 5, then 20 mg/L | 14 | 8.81% | - | - | - |
| *SAL1-KO* | 210 | 0.8 | 2 h | 1 h | 5, then 20 mg/L | 8 | 3.33% | 8 | 2 | 25.00% |
| *PDS-KO* | 212 | 0.8 | 2 h | 1 h | 5, then 20 mg/L | 10 | 4.72% | 10 | 2 | 20.00% |
| *CRY4-KO* | 526 | 0.8 | 2 h | 1 h | 5, then 20 mg/L | 28 | 5.32% | - | - | - |
| *YSL-KO* | 468 | 0.8 | 2 h | 1 h | 5, then 20 mg/L | 21 | 4.49% | - | - | - |

The pRGEB32-CrActin (Figure 2A), and pCAMBIA1300 (Figure 2B) were employed for gene editing and overexpression *C. richardii*, respectively. The transformation construct used for stable overexpression transformation was pCAMBIA1300-2×35S, which carries the *hygromycin phosphotransferase* (*HPT*) gene for selection of positive transgenic plants. After 72 h of co-incubation with *Agrobacteria*, transformed gametophytes were selected on MS media supplemented with 100 mg/L cefotaxime and 5 mg/L hygromycin to kill the *Agrobacteria* and select the transformants, respectively (Figure 3).

**Figure 2.**
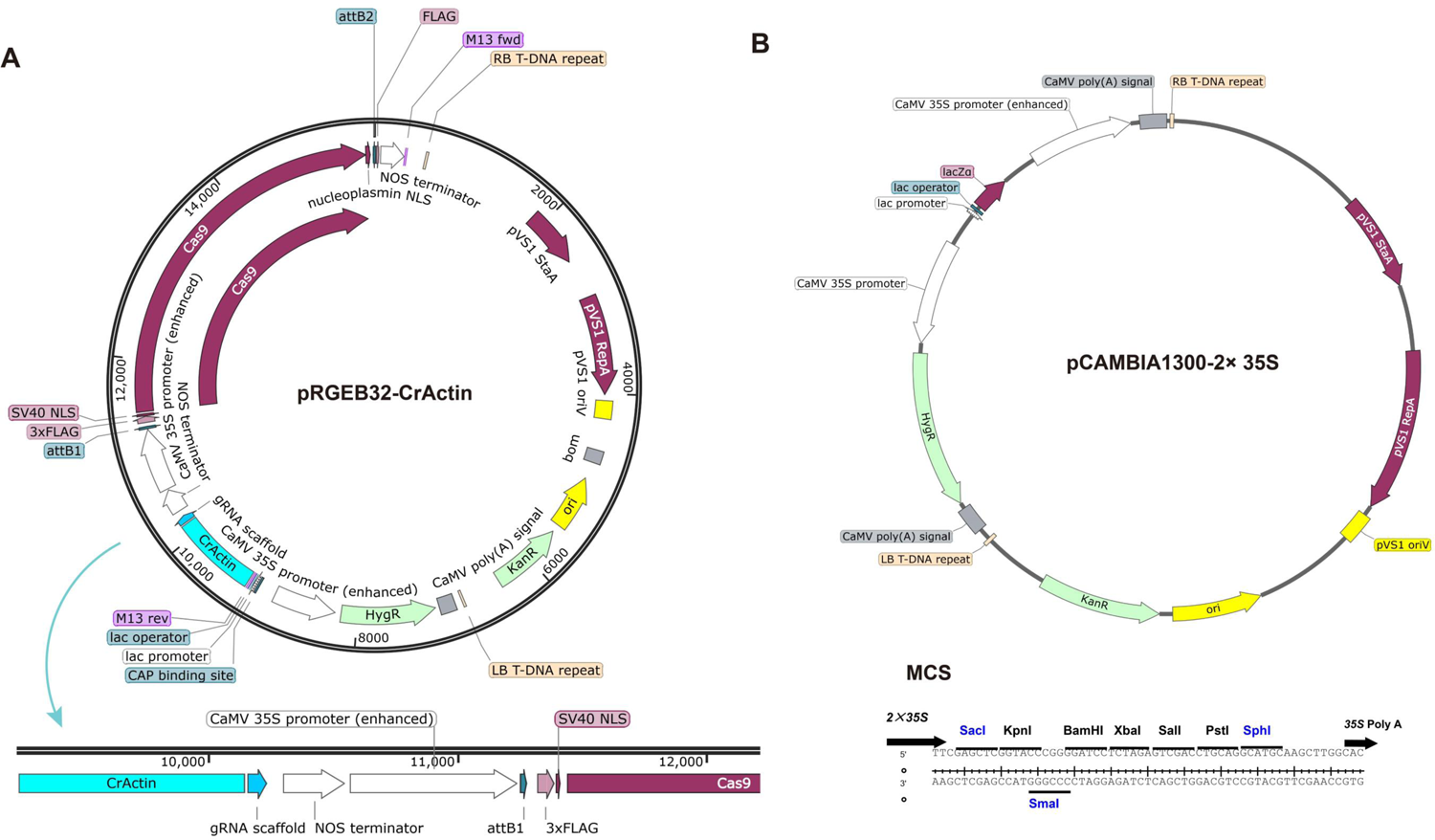
Plasmid information of CRISPR/Cas9 and overexpression for genetic transformation in *C. richardii*. CRISPR/Cas9 (A) and overexpression (B). A 916 bp fragment, located at the upstream of the *CrActin* was isolated and considered as the putative promoter, which was instead of the OsU3 promoter in pRGEB32 to drive the expression cassettes of sgRNA. The native maize ubiquitin promoter (ZmUbi) promoter in the original construct pRGEB32 was replaced by the enhanced 35S promoter, which was designated as pRGEB32-CrActin. The pRGEB32-CrActin and pCAMBIA1300 were employed for gene editing and overexpression *C. richardii*, respectively.

**Figure 3.**
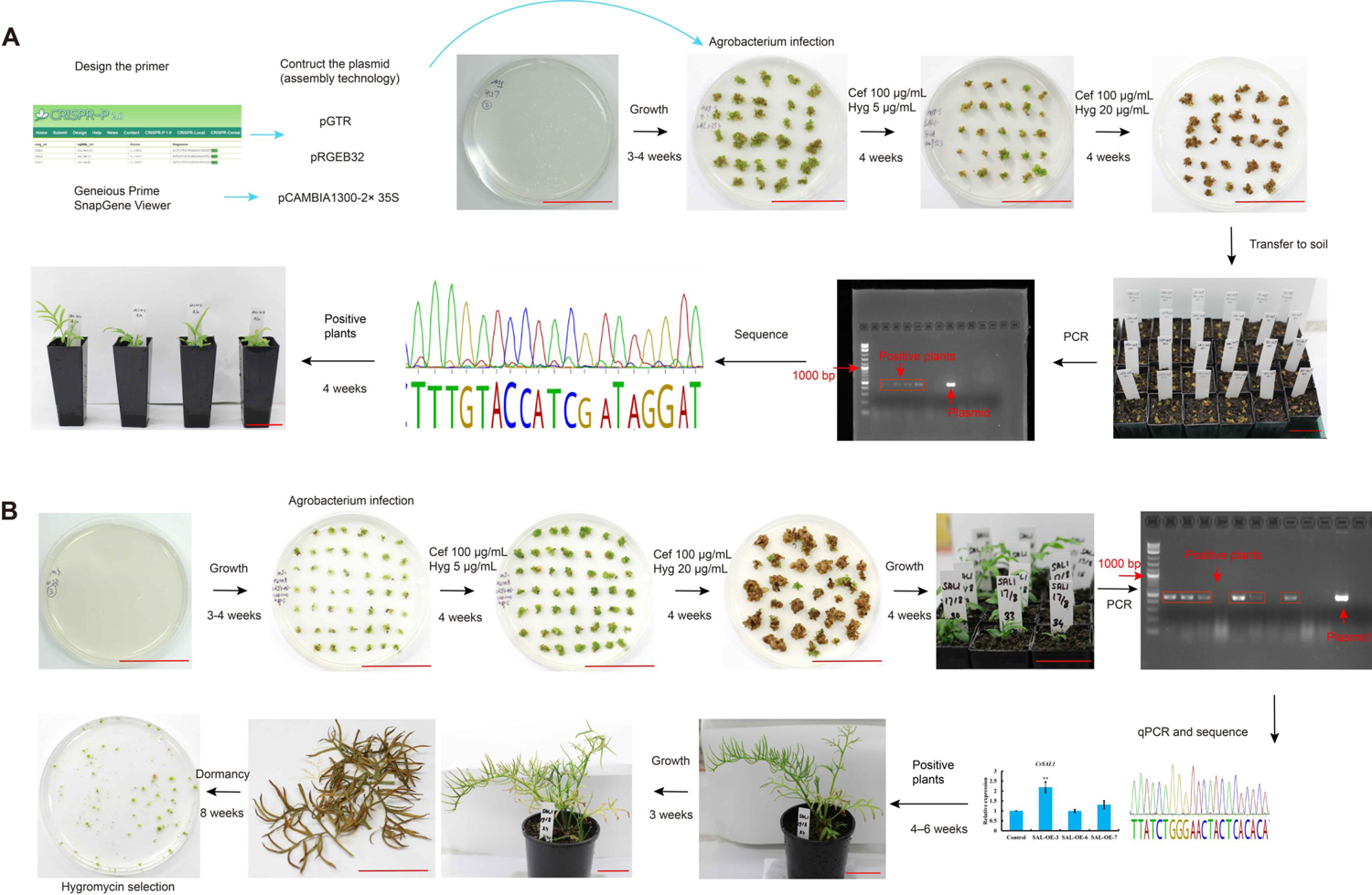
The workflow of gene editing (A) and gene overexpression (B) in *C. richardii*. Gene editing (A) and gene overexpression (B), bars = 5 cm. The sgRNAs were designed through CRISPR-P 2.0 (http://crispr.hzau.edu.cn/cgi-bin/CRISPR2/SCORE). Overexpression and CRISPR/cas9 constructs were generated utilizing the assembly technology. After *Agrobacterium* infection, gametophytes were grown at MS media with 5 mg/L of hygromycin and 100 mg/L of cefuroxime for 30 days. Then, the sporophytes were transferred to MS media supplemented with 100 mg/L cefotaxime and 20 mg/L hygromycin for another 30 days. The regeneration sporophytes were then transplanted to pots containing a premium potting mix for PCR and qPCR analysis.

We found that the gametophytes are unable to reproduce and survive for long periods under the suggested MS media with 20 mg/L hygromycin. In order to increase the regeneration and survival rate of the transformed gametophytes, we assayed a range of hygromycin concentrations and selected 5 mg/L (Supplemental Figure S3A, S3B), resulting in more regenerated gametophytes with normal morphology and reproduction (Figure 3). The sporophytes were then transferred to MS media supplemented with 100 mg/L cefotaxime and 20 mg/L hygromycin for another 30 days. The highest regeneration rate for stable transformation was achieved by 2 h treatment with 1.5% (w/v) cellulase before *Agrobacterium* co-incubation. We observed that sporophyte survival rate was slightly increased by *Agrobacterium* co-incubation time with 1.5% cellulase for 1 h (Table 2). Therefore, a combination of digestion with 1.5% cellulase and selection with 100 mg/L cefotaxime and 5/20 mg/L hygromycin was employed in our experiments. Interestingly, regeneration rarely occurs in a 1:1 stoichiometry, and a cluster of diverse regenerated gametophytes were developed from a gametophyte inoculated with *Agrobacterium* (Figure 3B). The regenerated sporophytes were then transplanted to pots containing a premium potting mix for further analysis.

### Molecular analysis of transgenic C. richardii plants

Nearly 10% of treated gametophytes survived on MS media supplemented with 20 mg/L hygromycin (Figure 3B). We obtain 87 *CrSAL1* overexpressed plants survived under hygromycin selection, but half of the plants failed to develop normally and complete the life cycle (Supplemental Figure S4A). Positive transgenic plants were screened by PCR with a 456-bp PCR product using the DNA as template and hygromycin primers targeting to the hygromycin gene (Supplemental Figure S4B). In total, we obtained and verified 15 transgenic *C. richardii* individuals with relatively higher expression of *CrSAL1* (Supplemental Figure S4). The transformation efficiency was calculated according to the number of successfully developed transgenic sporophytes divided by the total gametophytes used in transformation and multiplied by 100 (Bui et al., 2017), resulting in an efficiency ranging from 3.3% to 11.68% across those tested genes (Table 2).

### Screening of knockout lines of CrPDS and CrSAL mediated by CRIPSR/Cas9

After successful establishment of the *Agrobacterium*-mediated stable transformation method for overexpression gene of interest in *C. richardii* using gametophytes as the explant, the pipeline was employed to generate the gene editing lines with CRISPR/Cas9 system (Supplemental Figure S5) in *C. richardii* – the first of any fern species. Loss-of-function of *Phytoene desaturase* (*PDS*) leads to photobleaching phenotypes in varied plant species (Awasthi et al., 2021), which was widely employed as a visible marker in developing the protocol for knocking out of genes of interest (Ma et al., 2019). To introduce mutations into the *CrPDS*, two independent 20 bp sequences with NGG in their 3’-regions targeting were synthesized and inserted into the gRNA expression cassette of pRGEB32-CrActin vector. We obtained 18 *CrSAL1* and *CrPDS* CRIPSR/Cas9 plants through screening with hygromycin (Supplemental Figure S5). The positively transformed plants showed the expected photobleached leaf phenotype (Figure 4A). Sequence analysis determined that the editing efficiency of the *CrPDS* and *CrSAL1* target site in the transgenic plants was ranged from 20% to 25%, although the transformation efficiency of gene editing ranged from 3.33% to 4.72%. Both of replacement and deletion could be found in the mutant lines (Figure 4B, 4D). These results suggest that the pRGEB32-CrActin we generated in this study could be employed for editing genes of interest in *C. richardii* (Table 2).

**Figure 4.**
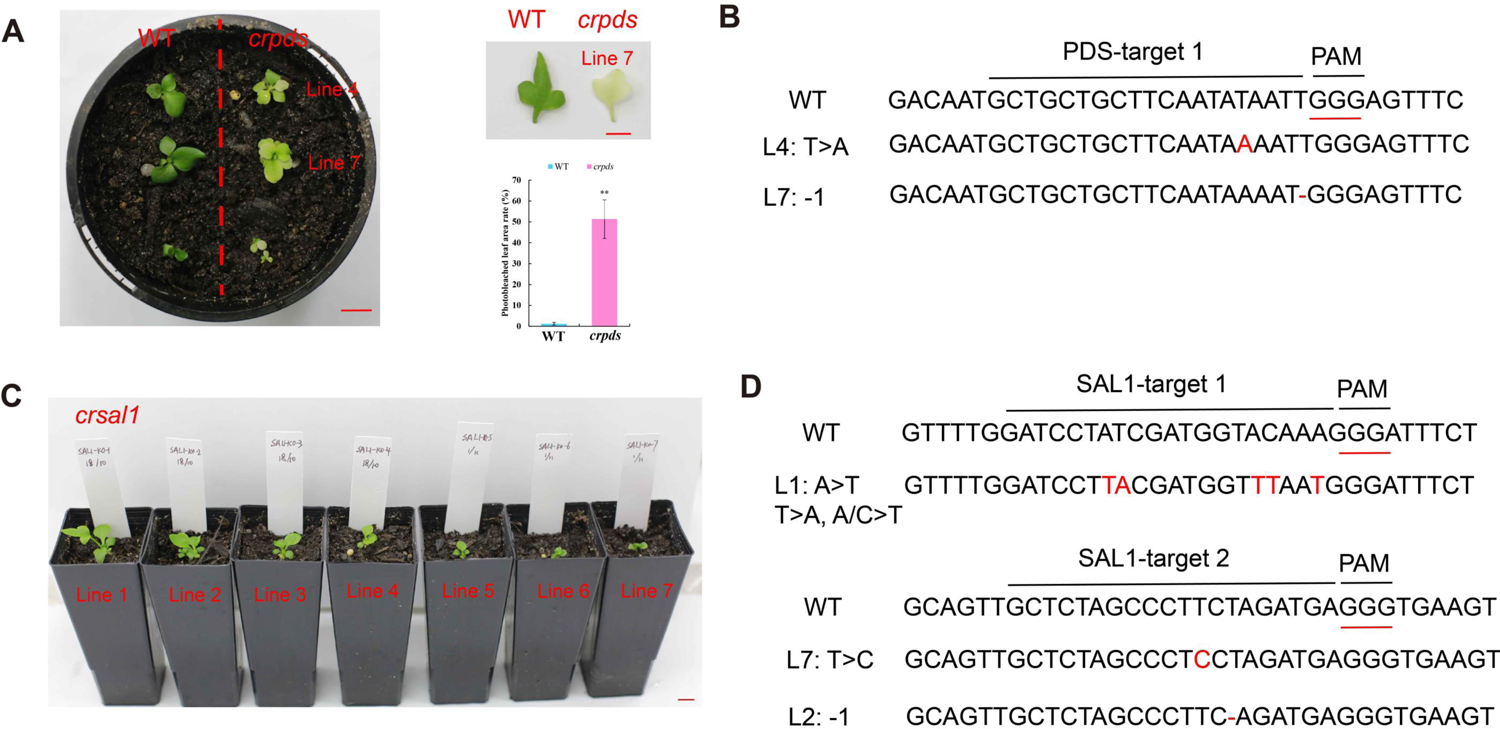
Phenotype and sequences of the editing types of *CrPDS* and *CrSAL1* transgenic plants. Phenotypes of WT and transgenic plants (*CrPDS*) (A), bars= 1 cm. Sanger sequencing of the editing types in *CrPDS* transgenic plants (B). Phenotypes of *crsal1* plants. Sanger sequencing of the editing types in *CrSAL1* transgenic plants (D).

### Physiological evaluation of SAL1 overexpression and knockout *C. richardii* plants

The subcellular localization of GFP fusion construct in the tobacco epidermis showed that GFP alone was found in the nuclei, cytoplasm, and membranes. However, we found GFP fluorescence of CrSAL1 overlaps with the chloroplast fluorescence, implying that the CrSAL1 protein is localized at the chloroplast and potentially in the cytosol (Figure 5C). The results indicate that CrSAL1 may function in chloroplast retrograde signaling and stomatal regulation similar to those seen in *A. thaliana* (Xiong et al., 2001; Estavillo et al., 2011).

**Figure 5.**
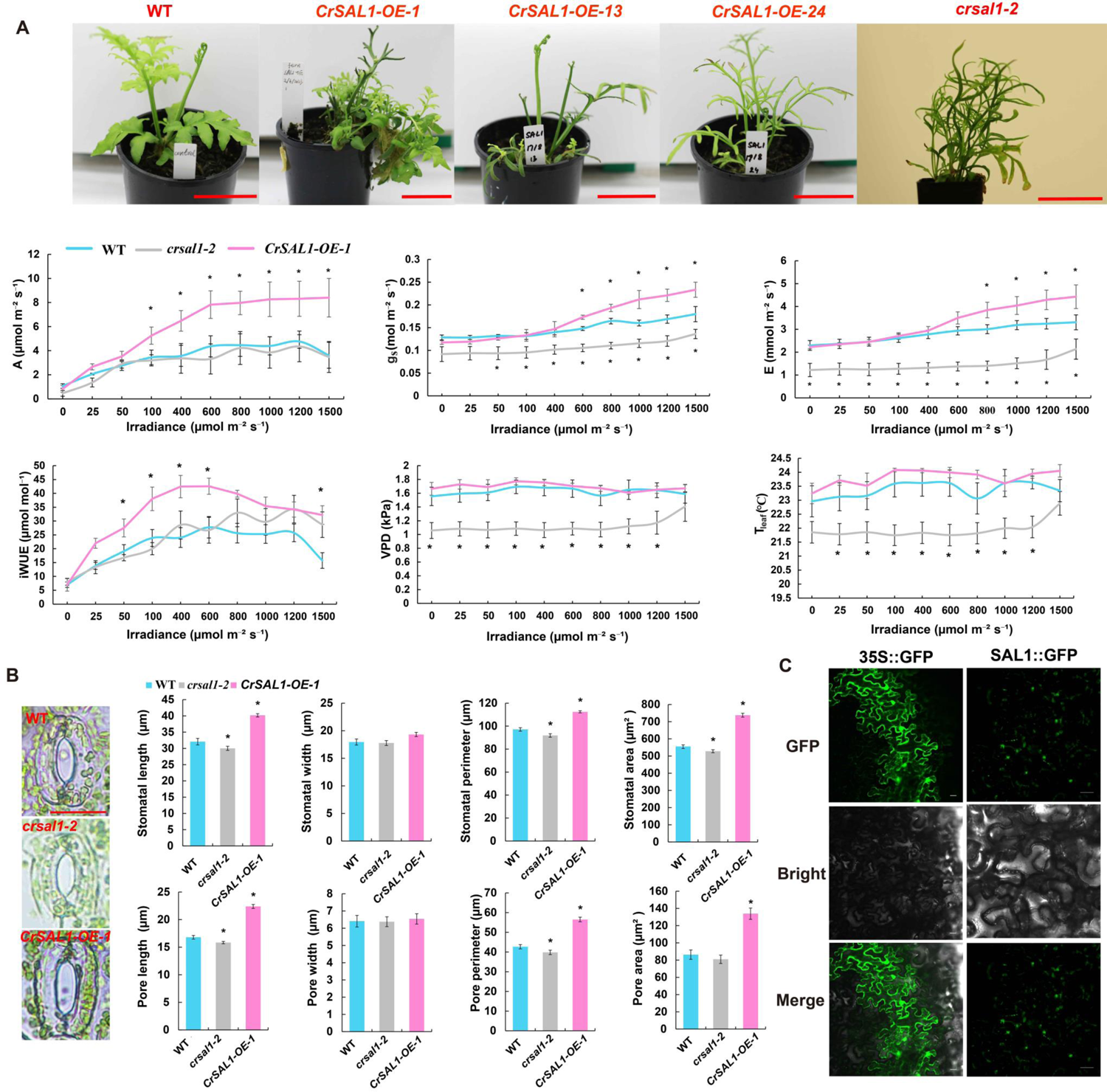
Photosynthesis and stomatal traits of *CrSAL1* gene editing and overexpression lines. Phenotype and gas exchange parameters of WT, *SAL1-OE*, and *crsal1* plants (A), bars= 5 cm. Net CO_2_ assimilation (*A*), leaf transpiration rate (*E*), stomatal conductance (*g_s_*), intrinsic water use efficiency (*iWUE*), vapor pressure deficit (*VPD*), and leaf temperature (*T_leaf_*) of transgenic and WT plants (n=6). Stomatal parameters of transgenic and WT plants (B), bar= 20 μm. Subcellular localization patterns of CrSAL1 in tobacco leaves (C), bars= 20 μm. Values are means of three biological replicates ± SE with 20–30 stomata. Asterisks indicate significant differences compared with the WT plants (\**P* < 0.05).

We overexpressed *CrSAL1* in *C. richardii* and obtained 15 individuals with relatively higher expression of *CrSAL1*, but only four individuals (Line 1, 13, 21 and 24) completed the life cycle (Figure 5A). Overexpression *CrSAL1-OE-1* (Line 1) in *C. richardii* significantly increased the net CO_2_ assimilation (*A*), leaf transpiration rate (*E*), and stomatal conductance (*g_s_*) under high light intensity compared to the WT across light intensity from 0 to 1500 μmol m^−2^ s^−1^. Interestingly, the *crsal1-2* CRIPSR/Cas9 knockout mutants displayed significantly lower *g_s_, E*, vapor pressure deficit (*VPD*), and leaf temperature (*T_leaf_*) compared to the WT (Figure 5A).

Stomata are essential for plants to respond to environmental conditions (Hetherington and Woodward, 2003; Chen et al., 2017; Jiang et al., 2024). In the control conditions, the *CrSAL1-OE-1* transgenic plants exhibited larger length, area, and perimeter of both stomata and stomatal pores compared to the WT plants (Figure 5B). Moreover, stomatal length, stomatal perimeter, and stomatal area in the *CrSAL1-OE-1* lines were significantly increased, on average, by 25.3%, 15.8%, and 30.4%, respectively. The mean pore length, pore perimeter, and pore area of *CrSAL1-OE-1* were increased by 33.3%, 32.4%, and 55.0%, respectively. In contrast, *crsal1-2* knockout mutants showed a slight decrease in the length and perimeter of stomata and stomatal pore compared to the WT (Figure 5B).

*CrSAL1-OE-1* plants also exhibited high ROS levels in guard cells under the control conditions. The total ROS level of *crsal1-2* plants was significantly lower than that of WT in the control conditions (Figure 6A), similar to the results of previous studies analyzing mutants of *SAL1* gene such as *altered ascorbate peroxidase 2* (*APX2*) *expression 8* (*alx8*) and *onset of leaf death 101* (*old101*) in *A. thaliana* (Estavillo et al., 2011; Shirzadian-Khorramabad et al., 2022). SAL1 was reported to be important for ABA signaling in response to environmental conditions (Pornsiriwong et al., 2017; Zhao et al., 2019). Thus, we also performed the stomatal assay with ABA treatment in the WT and transgenic plants. Interestingly, *crsal1-2* mutant displayed ABA-sensitive stomatal phenotype (Figure 6C, 6D), which is consistent with the previous study that *sal1-8* (Pornsiriwong et al., 2017) and *fry1* (Xiong et al., 2001) were more sensitive to ABA in *A. thaliana*, implying the potentially conserved molecular function of *SAL1* in stomatal regulation in different plants. Furthermore, ABA treatment increased the ROS level of guard cell in WT, *crsal1-2, CrSAL1-OE-1* plants (Figure 6B), leading to stomatal closure. In summary, we demonstrated for the first time on the gene editing in *C. richardii* by editing four important genes and anlayzed the function of *CrSAL1*.

**Figure 6.**
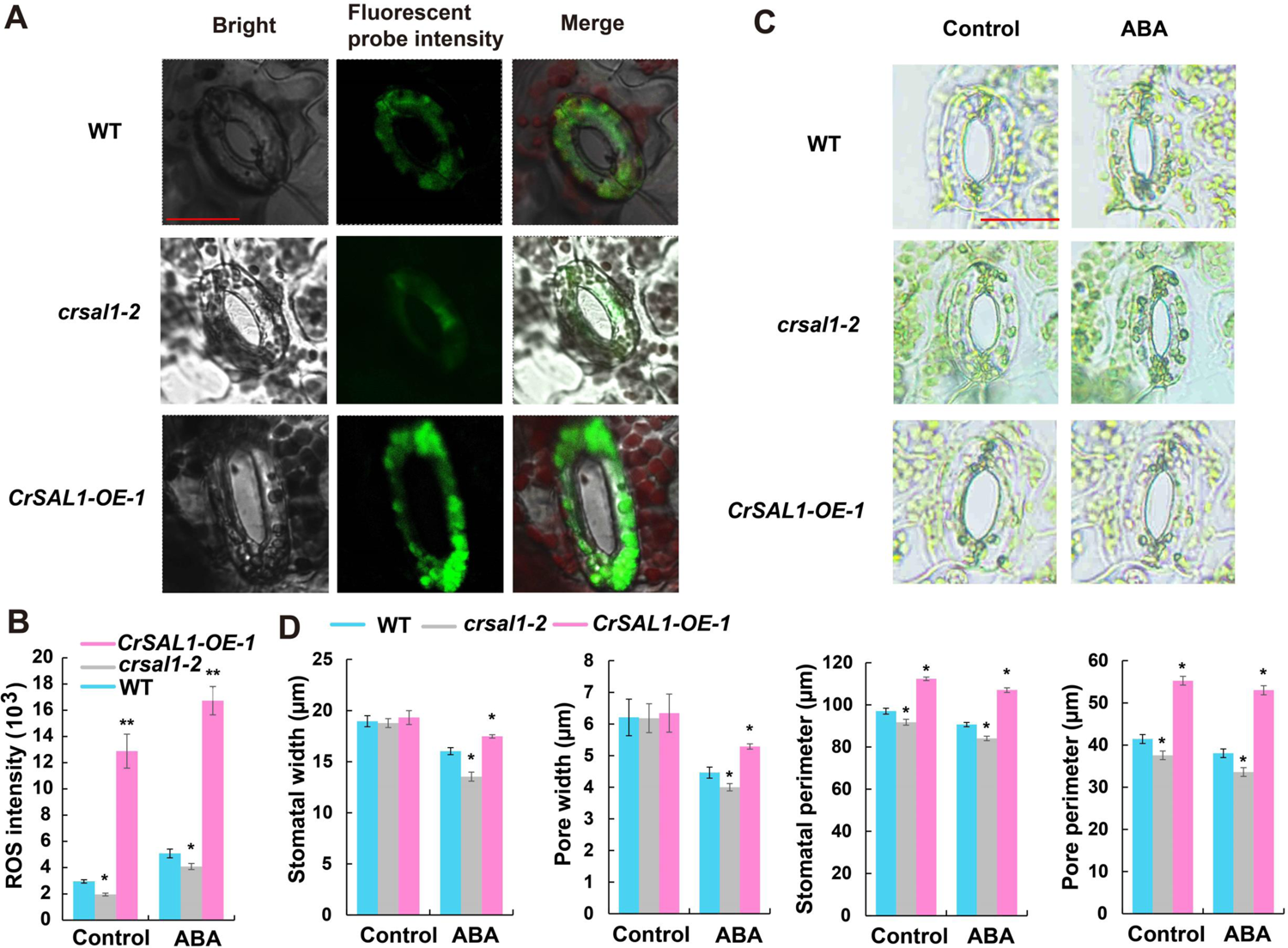
Effects of gene editing and overexpression *CrSAL1* on reactive oxygen species (ROS) and ABA response of fern plants. Confocal images and fluorescent probe intensity of ROS in guard cells of the *SAL1* CRISPR/Cas9 and overexpression plants (A), bar= 20 μm. ABA induced stomatal close in transgenic and WT plants (C), bar=20 μm. ROS intensity (B) and stomatal traits (D) and of *C. richardii* in control and 50 μΜ ABA treatment for 60 min. Values are means of three biological replicates ± SE with 20–30 stomata. Asterisks indicate significant differences compared with the WT plants (\**P* < 0.05, \*\**P* < 0.01).

## Discussion

### First gene editing for gene functional verification in a fern

CRISPR/Cas genome editing has been applied to a variety of plant species to enhance disease resistance and abiotic stress tolerance (Deng et al., 2022). In the past ten years, there were 9,000 publications on topics relevant to plant CRISPR on Web of Science (https://www.webofscience.com/). However, there have been no studies on the use of CRISPR/Cas9 in ferns (Frangedakis et al., 2023). In this study, we established an efficient gene editing method for the transformation of *C. richardii*.

We successfully overexpressed *CrSAL1* and other genes in *C. richardii* gametophytes by adjusting the hygromycin concentration (Bui et al., 2015), OD value of *Agrobacterium*, age of gametophytes, and enzyme treatment time of gametophytes and co-cultivation with *Agrobacterium* (Table 2, Figure 3). This optimized protocol enabled us to establish stable *Agrobacterium*-mediated CRISPR/Cas9 transformation in *C. richardii*. Due to the low expression of *CrU6* genes (Supplemental Figure S1A) and the low efficiency of ZmUbi in *C. richardii*, the OsU3 and ZmUbi promoter of pRGEB32 plasmid were replaced by the promoter of *CrActin* and enhanced 35S, respectively. This system can edit genes with high efficiency in *C. richardii* based on the success with *CrSAL1, CrPDS* and other genes. In most of the CRISPR/Cas9 constructs, the RNA polymerase III-type 3 - U3 or U6 promoters are employed for expression of sgRNA in monocots, eudicots, gymnosperms, and bryophytes (Kor et al., 2022). Although we did not use CrU6 promoters due its low expression, we hypothesize that they can have potential applications in the genome editing of ferns. The ZmUbi had been successfully used for generating the RNAi plants of *C. richardii* (Plackett et al., 2018), but it requires further investigation on whether it can be used for fern gene editing or not.

In the future, direct transformation of gametophytes for gene functions in apogamy (Bui et al., 2018) may provide a clue to the evolution of asexual reproduction in land plants, permitting comparison of fern apogamy to somatic embryogenesis and apomixis in angiosperms (Kinosian and Wolf, 2022). Therefore, once the current gene editing method of *C. richardii* is applicable to many other fern species, we can study key biological aspects such as the role of duplicate genes as well as physiological features and the evolution of stress tolerance in ferns at the molecular level using gene editing. Despite the great potential, several issues still limit the efficiency of CRISPR/Cas9 as a tool for mitigating plant stresses (Deng et al., 2022). For instance, the inactivation of some genes through gene editing often results in disease resistance, but is also associated with pleiotropic effects such as inhibition of plant growth, phenotypic abnormalities and increased susceptibility to abiotic stress and other pathogens (Ma et al., 2018). Abiotic stress tolerance usually depends on complex mechanisms controlled by multiple genes (Adem et al., 2020; Shabala et al., 2020; Tripathi et al., 2020; Wang et al., 2023), implicating the need to develop multiplex CRISPR-based approaches for ferns.

### Advantages of using gametophytes in the transformation of ferns

Bryophytes, ferns and lycophytes rely on free-living gametophytes for reproduction (Fouracre and Harrison, 2022). Unlike mosses and liverworts whose dominant generation is the gametophyte (Frangedakis et al., 2023), the dominant generation in ferns is the sporophyte. The spores of ferns are shed by the sporophytes and develop into free-living gametophytes (Bui et al., 2018). This life cycle of ferns provides an opportunity to use gametophytes as targets for transgenesis (Kinosian and Wolf, 2022). This is in stark contrast to the transformation protocol for angiosperm species, where the immature embryo, callus, flowers and protoplasts are usually used for efficient stable transformation (Altpeter et al., 2016). The advantages of using gametophytes are relatively simple and reproducible using large quantity of spores (Bui et al., 2018), which are fast to germinate, easy to manage, and quick to grow on solid medium compared to laborious embryo separation and callus induction needed for genetic transformation of many angiosperms (Ishizaki et al., 2016).

RNAi was made possible through direct uptake of dsRNA into germinating spores of *C. richardii* (Stout et al., 2003) and *Marsilea vestita* (Klink and Wolniak, 2001). Particle bombardment of DNA constructs into gametophytes has also been demonstrated in *C. richardii* (Rutherford et al., 2004) and *Adiantum capillus-veneris* (Kawai-Toyooka et al., 2004) (Table 1). Transgenesis in ferns was demonstrated in *C. thalictroides* and *P. vittata* with five-day-old germinating spores and 15-day-old gametophytes by *Agrobacterium*-mediated transformation and particle bombardment transformation, respectively (Muthukumar et al., 2013).

A tractable particle bombardment transgenesis system using sporophytes has been established in *C. thalictroides* and *C. richardii* (Plackett et al., 2014; Plackett et al., 2015). Callus tissues were induced from young sporophytes, and then bombarded with a GUS reporter and hygromycin selection (Plackett et al., 2014). This method requires callus induction similar to transformation protocols of angiosperms followed by sporophyte regeneration. Here, we optimized the enzyme treatment time, OD value of *Agrobacterium*, the suitable concentrations of hygromycin selection, and planting density in Petri dishes, achieving a higher transformation efficiency close to 10% in overexpression of *CrSAL1* (Table 1, Figure 3). The high transformation efficiency will benefit better understand the function of important genes in the biology, evolutionary, and future agricultural and medicinal applications of ferns.

### Conserved evolution and functional divergence of SAL genes family

Plant *SAL1s* have been extensively reported to be involved in phytohormones (Ishiga et al., 2017) (e.g., ABA, salicylic acid, jasmonic acid, and auxin) and stresses (Jia et al., 2019) such as *Fusarium graminearum* (Yu et al., 2015), salt (Chen et al., 2011), drought (Abdallah et al., 2022), cold (Shen et al., 2023), high light (Estavillo et al., 2011), oxidative stress (Chan et al., 2016), and cadmium (Xi et al., 2016). Due to its distinct effects on different cellular processes, the underlying molecular mechanisms of *SAL1* in stress responses appears to be complex (Jia et al., 2019). In *A. thaliana*, there are four SALs (AT5G63980, AT5G64000, AT5G63990, AT5G09290) and two homologs [inositol monophosphatase, AT5G54390 (Arabidopsis Halotolerance 2-like, AHL) and AT4G05090] (Shin et al., 2019). AtSAL1 plays a negative role in stress response pathways that are predominantly ABA-dependent and ABA-independent (Wilson et al., 2009).

The *C. richardii* genome contains one inositol-1,4-bisphosphate 1-phosphatase CrHAL2/CrSAL2 (Ceric.01G129600) and PAP-specific phosphatase CrHAL2-like (Ceric.11G097700), which shows 32% and 44% identity to 3’(2’), 5’-bisphosphate nucleotidase CrSAL1 (Ceric.25G052000), respectively. In this study, *crsal1-2* mutant displayed ABA-sensitive stomatal phenotype, which is in accordance with *fry1* (Xiong et al., 2001) and *sal1-8* (Pornsiriwong et al., 2017) that were more sensitive to ABA in *A. thaliana* compared to the WT. In addition, overexpressed *CrSAL1-1* plants exhibited reduced response to ABA-induced stomatal closure, which is in agreement with previous report that ectopic expression of soybean *GmSAL1* in *A. thaliana* decreased the ABA-induced stomatal closure (Ku et al., 2013). *A. thaliana alx8* also showed low *A* and *g_s_* (Rossel et al., 2005) and *A. thaliana old101* mutants of maintained lower ROS levels (Shirzadian-Khorramabad et al., 2022). Interestingly, *crsal1-2* showed significantly lower ROS production in the guard cell and decreased photosynthetic parameters (e.g. *A, g_s_, VPD, T_leaf_*) (Figures 5 and 6), indicating the functional similarity of *SAL1s* in the two species.

Our previous study showed that SAL1 and its chloroplast transit peptides were conserved in chlorophyte algae and land plants (Zhao et al., 2019). 197 *SAL* genes in 53 *Chlorophyta* and *Embryophyta* species were identified (Supplemental Figure S6) through PLAZA platform (Van Bel et al., 2022) with 27% and 17% of block and tandem within this gene family. The *SAL* gene family was greatly expanded in monocots (e.g., *T. aestivum, Phyllostachys edulis*) and eudicots (e.g., *Glycine max, Brassica napus*), but not in bryophytes and ferns. Gene expression profiles of *SALs* showed that some genes are specifically expressed in the reproductive organs, leaf, and root (Proost and Mutwil, 2018). Interestingly, *AtSAL1* showed high expression in many organs such as root, stem, leaf, flower, seed, reproductive organs, and meristem (Table S2). *AtSAL2* was preferentially expressed in the leaf, while *AtSAL4* displayed specific expression in root, implying their different roles in these tissues. Drought induced the expression of *Zm00001e039578_P001* (*GRMZM2G152757, SAL1*) in maize (Kim et al., 2021), which was also involved in photoperiod at vegetative-tasseling stage (Wang et al., 2017) and osmotic stress (Yu et al., 2018). Interestingly, red fluorescence of RFP-SAL (Pp3c3_21240V3.1) was observed in the cytosol of moss *Physcomitrella patens* cells (Cross et al., 2017), implying the diverse biological functions of SALs. Expression analyzes demonstrated that some *SAL* genes function in leaf and root of gymnosperms and lycophytes and others are important for the reproductive organs of angiosperms, illustrating that neofunctionalization of *SAL* genes might coincide with the emergence of expansion in angiosperms. However, the study of SALs mainly focused on the *A. thaliana* (Jia et al., 2019). Thus, investigations of the molecular mechanisms of SALs through gene editing are important for enhancing abiotic stress tolerance in crops and addressing key evolutionary biology questions in important early divergent plant lineages such as ferns.

## Materials and Methods

### Plant materials and growth conditions

The *C. richardii* genotype Hn-n with a fully sequenced and assembled genome (Marchant et al., 2022) was used in our study. Plants were grown in a GEN 1000 (CONVIRON, Manitoba, Canada) at 16 h of light/8 h of dark, 28°C, 80% relative humidity, and fluence of 100 μmol m^-2^ s^-1^. Gametophytes were grown with 1.5% (w/v) of 1× Murashige and Skoog (MS) in agar medium at pH of 5.9 (Plackett et al., 2015). Spores were sterilized by incubating for 5 min in sodium hypochlorite solution [1% (v/v) chlorine], which was subsequently removed by three sequential rinses with sterile distilled water at 23°C. Spores were then imbibed in distilled water and incubated for 3 days in darkness before sowing (Plackett et al., 2014; Withers et al., 2023). The spores were imbibed in 1 mL sterile water in the Petri dish, which was sealed with foil and incubated at 28°C for 7 days and germinated. One-month-old gametophytes can be used for the transformation of *Agrobacterium*.

### Agrobacterium tumefaciens-mediated transformation of gametophytes

Stable genetic transformation of *C. richardii* plants was performed as described previously with modification (Bui et al., 2015; Bui et al., 2017). More details can be found in Supplementary Materials and Methods. Overexpression and CRISPR/cas9 constructs were generated utilizing the assembly technology (Bai et al., 2020). Briefly, the PCR products of full-length coding sequences (CDS) were cloned into the vector pJET1.2/blunt using CloneJET PCR Cloning Kit (Thermo Fisher Scientific, Waltham, MA USA) (Awasthi et al., 2021), and then transformed into DH5α competent cells (Life Technologies, Waltham, MA USA). Plasmid purification was performed with a GeneJET Plasmid Miniprep Kit (Thermo Fisher Scientific, Waltham, MA USA) (Lorenzo et al., 2023) and the resulting plasmid DNAs were validated by sequencing. The correct sequence was introduced into the destination vectors pCAMBIA1300-2× 35S [enhanced cauliflower mosaic virus (CaMV) 35S promoter] at the restriction enzyme sites *BamHI* and *PstI* (New England BioLabs, Ipswich, MA, USA). The sgRNAs were designed through CRISPR-P 2.0 (http://crispr.hzau.edu.cn/cgi-bin/CRISPR2/SCORE) (Liu et al., 2017). To generate CRISPR/Cas9 plasmid, fragments containing tRNA-sgRNA1 fusion and gRNA-tRNA-sgRNA2 fusion were obtained through pGTR as a template (Xie et al., 2015; Wang et al., 2018). The PCR products were then cloned into pRGEB32-CrActin vector at the restriction enzyme site *BsaI* (Fu et al., 2022; Kuang et al., 2022). All constructs were introduced into the *Agrobacterium tumefaciens* strain GV3101.

After *Agrobacterium* infection, gametophytes were grown in MS media with 5 mg/L of hygromycin and 100 mg/L of cefuroxime for 30 days. Then, the sporophytes were transferred to new MS media containing 20 mg/L of hygromycin and 100 mg/L of cefuroxime for 30 days. T1 sporophytes grown without hygromycin selection and transgenic individuals were subsequently identified by hygromycin selection on MS media (Plackett et al., 2015). Sporophytes were then transplanted to pots containing a premium potting mix (Scotts Osmocote, Bella Vista, Australia) with the cover to keep high humidity. The plants were watered and fertilized fortnightly with a nutrient solution at the 0.5 g/L (Thrive Soluble Fertilizer, Yates, Padstow NSW, Australia).

### qPCR analysis of transgenic plants

For expression analysis of *CrSAL1*, total RNA was extracted from infertile leaves through RNeasy Plant Mini Kit (QIAGEN) (Cai et al., 2017; Cai et al., 2021). The cDNA synthesis was performed by QuantiTect Reverse Transcription Kit (QIAGEN) and the synthesized cDNA was diluted five times before Quantitative real-time PCR (qPCR) experiments. The qPCR was conducted for three biological replicates using a QuantiNova SYBR Green PCR Kit (QIAGEN) on a LightCycler 96 Real-Time PCR System (CFX Connect) (Jiang et al., 2020). Expression levels were normalized against the *CrACTIN* reference gene (Plackett et al., 2018). The relative expression levels of genes were performed from cycle threshold values by 2^−ΔΔCt^ procedure (Feng et al., 2020; Jiang et al., 2022). All primers were designed using Primer Premier 6.0 (PREMIER Biosoft, San Francisco, CA, USA) or SnapGene Viewer (GSL Biotech LLC, Boston, MA, USA) in this study (Supplemental Table S1).

### Subcellular localization

Subcellular localization of CrSAL1 was performed according to the previous study (Feng et al., 2020). The coding regions of CrSAL1 were amplified and cloned into pCAMBIA1300-GFP (Fu et al., 2022) by the restriction enzyme site *KpnI* and *XbaI*. The resulting plasmids were transferred into *A. tumefaciens* strain GV3101. *A. tumefaciens* harboring the vector was grown overnight in Luria broth (LB) medium containing 25 mg/L of Rifampin and 50 mg/L of Kanamycin (Jiang et al., 2022). After centrifugation, *A. tumefaciens* was resuspended through the infiltration buffer [10 mM 2-(N-morpholino) ethanesulfonic acid (MES)-KOH (pH 5.7), 10 mM MgCl_2_, 100 μM acetosyringone (AS)] to achieve OD600 = 0.8. The suspension was infiltrated into the abaxial air spaces of 4-week-old *Nicotiana benthamiana* leaves using a 1-mL syringe without a needle to transiently express (Feng et al., 2020). Green fluorescent protein (GFP) fluorescence was detected through using a confocal microscopy (SP5, Leica Microsystems GmbH, Wetzlar, Germany) (Deng et al., 2021).

### Measurement of reactive oxygen species (ROS)

The production of ROS in guard cells of *CrSAL1* transgenic and WT plants was measured using a fluorescent indicator 2’,7’-dichlorodihydrofluorescein diacetate (CM-H_2_DCFDA, Life Technologies, Waltham, MA USA) (Cai et al., 2017). epidermal peels were incubated with an opening buffer [10 mM KCl and 5 mM MES at pH 6.1 with Ca(OH)_2_] for 30 mins for stomatal assays epidermal peels The samples were then loading with 20 µM CM-H_2_DCFDA for 30 min in the dark, followed by a 5 min rinse with a measuring buffer [50 mM KCl and 10 mM MES at pH 6.1 with NaOH] to remove excess dye (Cai et al., 2021). The epidermal peels were then incubated in the measuring buffer for confocal microscopy imaging with excitation at 488 nm and emission at 510–540 nm (SP5, Leica Microsystems GmbH, Wetzlar, Germany).

### Gas exchange measurements

Gas exchange measurements were measured on the *C. richardii* fully expanded infertile leaf by LI-6400 infrared gas analyzer (LI-COR, USA) (Liu et al., 2017; Qiu et al., 2023). The parameters are net CO_2_ assimilation (*A*), stomatal conductance (*g_s_*), leaf transpiration rate (*E*), vapor pressure deficit (*VPD*), and leaf temperature (*T_leaf_*). The intrinsic water use efficiency (*iWUE*) calculation is the ratio of *A* to *g_s_*. Leaf chamber conditions were maintained at a flow rate of 500 mol s^−1^, 70% relative humidity and 400 ppm reference CO_2_. Irradiance levels were set at 0, 20, 50, 100, 200, 300, 500, 800, 1000, and 1500 μmol m^−2^ s^−1^ for light response curve measurement.

### Stomatal assay

Stomatal assay was determined from the abaxial surface of the fully expanded and mature leaves as described in our previous work (O’Carrigan et al., 2014; Liu et al., 2017; Plackett et al., 2021). For these measurements, fully expanded infertile leaves were removed from the chamber and placed in Petri dishes on tissue paper soaked in opening buffer. The lower leaf epidermis was quickly peeled off and placed it on slides with the opening buffer. Stomatal morphology was calculated from the leaf epidermis through a light microscopy and imaging system (Nikon, Tokyo, Japan). Treatment was applied as ABA (50 μM) measured for another 60 min. The pictures were imported into the ImageJ software for the analysis of multiple parameters. Stomatal area (total stomatal area), stomatal perimeter (total length of the stomatal outer border), stomatal length (top to bottom of the stomatal), stomatal width (left to right of the stomatal), pore area (total pore area), pore perimeter (total length of the pore outer border), pore length (top to bottom of the pore), and pore width (left to right of the pore) were recorded (Pan et al., 2022). There were 20–30 stomata with three biological replicates for each treatment and genotype.

### Statistical analysis

Data were shown as means with standard errors of three biological replicates. The SPSS 26.0 software (IBM, USA) was employed to perform the analysis of variance (ANOVA) and means were compared by Duncan’s multiple range tests.

## Funding

Z-HC is funded by the Australian Research Council (FT210100366), Horticulture Innovation Australia (LP18000), CRC Future Food Systems (P2-016, P2-018) and Grain Research and Development Corporation (WSU2303-001RTX). WJ was funded by the China Scholarship Council (202208420191). FZ and FD were funded by the National Natural Science Foundation of China (32272053, 32370285, and 32170276).

## Acknowledgments

We thank Dr Yuanyuan Wang and Basanti Bhatta (Western Sydney University) and Dr Chenchen Zhao (University of Tasmania) for the discussion on the experiments. We thank Professor Kabin Xie (Huazhong Agriculture University) for providing pRGEB32 and pGTR vectors. We thank Professor Fei Chen (Hangzhou Normal University) for providing the vector of pCAMBIA1300-2× 35S, and Professor Dean Jiang (Zhejiang University) for providing the pCAMBIA1300-GFP vector.

## Author contributions

Z-HC planned and designed the research. WJ performed the experiments with research and technical support from MB. WJ and Z-HC analyzed the data and prepared the Figures and Tables. WJ, MB, and Z-HC analyzed the results and wrote the manuscript with support from CC, DY, TT, FD, and FZ. WJ, MB, DBM, PS, DS, FD, FZ and Z-HC conducted the editing of the manuscript. All authors read and approved the manuscript.

## Conflict of interest

The authors declare no conflict of interests.

## Supplementary data

The following materials are available in the online version of this article.

**Supplemental Figure S1.** Expression and alignment of sequences of U6 small nuclear RNA genes in diverse tissues of *C. richardii*.

**Supplemental Figure S2.** The sequences of the Actin/U6 promoter from *C. richardii* and 2× CaMV 35S promoter.

**Supplemental Figure S3.** Growth of gametophytes and sporophytes for testing hygromycin sensitivity and germination in *C. richardii*.

**Supplemental Figure S4.** Genotyping and phenotyping of *CrSAL1* overexpression *C. richardii* plants.

**Supplemental Figure S5.** Phenotype of *CrSAL1* (A) and *CrPDS* CRIPSR/Cas9 plants of *C. richardii* on the hygromycin selection medium.

**Supplemental Figure S6.** Tandem and block gene duplicate of *SAL* genes family in *Chlorophyta* and *Embryophyta*.

**Table S1.** List of primer sequences used in this study.

**Table S2.** Expression patterns of plant SALs in different tissues and evolutionarily important lineages of plants.

## References

Abdallah NA, Elsharawy H, Abulela HA, Thilmony R, Abdelhadi AA, Elarabi NI. Multiplex CRISPR/Cas9-mediated genome editing to address drought tolerance in wheat. GM Crops Food 2022: 1–17.

Adem GD, Chen G, Shabala L, Chen ZH, Shabala S. GORK channel: a master switch of plant metabolism? Trends Plant Sci. 2020: 25: 434–445.

Altpeter F, Springer NM, Bartley LE, Blechl A, Brutnell TP, Citovsky V, Conrad L, Gelvin SB, Jackson D, Kausch AP, et al. Advancing crop transformation in the era of genome editing. The Plant Cell 2016: 28: 1510–1520.

Awasthi P, Kocabek T, Mishra AK, Nath VS, Shrestha A, Matousek J. Establishment of CRISPR/Cas9 mediated targeted mutagenesis in hop (*Humulus lupulus*). Plant Physiol. Biochem. 2021: 160: 1–7.

Bai M, Yuan J, Kuang H, Gong P, Li S, Zhang Z, Liu B, Sun J, Yang M, Yang L, et al. Generation of a multiplex mutagenesis population via pooled CRISPR-Cas9 in soya bean. Plant Biotechnol. J. 2020: 18: 721–731.

Belshaw N, Grouneva I, Aram L, Gal A, Hopes A, Mock T. Efficient gene replacement by CRISPR/Cas-mediated homologous recombination in the model diatom Thalassiosira pseudonana. New Phytol. 2023: 238: 438–452.

Bui LT, Cordle AR, Irish EE, Cheng CL. Transient and stable transformation of Ceratopteris richardii gametophytes. BMC Res. Notes 2015: 8: 214.

Bui LT, Long H, Irish EE, Cordle AR, Cheng C-L (2018) The power of gametophyte transformation. In Current Advances in Fern Research, pp 271-284.

Bui LT, Pandzic D, Youngstrom CE, Wallace S, Irish EE, Szovenyi P, Cheng CL. A fern *AINTEGUMENTA* gene mirrors *BABY BOOM* in promoting apogamy in *Ceratopteris richardii*. Plant J. 2017: 90: 122–132.

Cai S, Chen G, Wang Y, Huang Y, Marchant DB, Wang Y, Yang Q, Dai F, Hills A, Franks PJ, et al. Evolutionary conservation of ABA signaling for stomatal closure. Plant Physiol. 2017: 174: 732–747.

Cai S, Huang Y, Chen F, Zhang X, Sessa E, Zhao C, Marchant DB, Xue D, Chen G, Dai F, et al. Evolution of rapid blue-light response linked to explosive diversification of ferns in angiosperm forests. New Phytol. 2021: 230: 1201–1213.

Cao H, Chai TT, Wang X, Morais-Braga MFB, Yang JH, Wong FC, Wang RB, Yao HK, Cao JG, Cornara L, et al. Phytochemicals from fern species: potential for medicine applications. Phytochem. Rev. 2017: 16: 379–440.

Cardi T, Murovec J, Bakhsh A, Boniecka J, Bruegmann T, Bull SE, Eeckhaut T, Fladung M, Galovic V, Linkiewicz A, et al. CRISPR/Cas-mediated plant genome editing: outstanding challenges a decade after implementation. Trends Plant Sci. 2023: 28: 1144–1165.

Čermák T, Curtin SJ, Gil-Humanes J, Čegan R, Kono TJY, Konečná E, Belanto JJ, Starker CG, Mathre JW, Greenstein RL, et al. A multipurpose toolkit to enable advanced genome engineering in plants. The Plant Cell 2017: 29: 1196–1217.

Chan KX, Mabbitt PD, Phua SY, Mueller JW, Nisar N, Gigolashvili T, Stroeher E, Grassl J, Arlt W, Estavillo GM, et al. Sensing and signaling of oxidative stress in chloroplasts by inactivation of the SAL1 phosphoadenosine phosphatase. Proc Natl Acad Sci U S A 2016: 113: E4567–4576.

Chen H, Zhang B, Hicks LM, Xiong L. A nucleotide metabolite controls stress-responsive gene expression and plant development. PLoS One 2011: 6: e26661.

Chen ZH. Unveiling novel genes in Fern genomes for the design of stress tolerant crops. Crop Des. 2022: 1: 100013.

Chen ZH, Chen G, Dai F, Wang Y, Hills A, Ruan YL, Zhang G, Franks PJ, Nevo E, Blatt MR. Molecular evolution of grass stomata. Trends Plant Sci. 2017: 22: 124–139.

Conway SJ, Di Stilio VS. An ontogenetic framework for functional studies in the model fern *Ceratopteris richardii*. Dev. Biol. 2020: 457: 20–29.

Cross LL, Paudyal R, Kamisugi Y, Berry A, Cuming AC, Baker A, Warriner SL. Towards designer organelles by subverting the peroxisomal import pathway. Nat. Commun. 2017: 8: 454.

Cui Y, Zhao J, Gao Y, Zhao R, Zhang J, Kong L. Efficient multi-sites genome editing and plant regeneration via somatic embryogenesis in *Picea glauca*. Front. Plant Sci. 2021: 12: 751891.

Deng F, Zeng F, Shen Q, Abbas A, Cheng J, Jiang W, Chen G, Shah AN, Holford P, Tanveer M, et al. Molecular evolution and functional modification of plant miRNAs with CRISPR. Trends Plant Sci. 2022: 27: 890–907.

Deng Z, Wu H, Jin T, Cai T, Jiang M, Wang M, Liang D. A sequential three-phase pathway constitutes tracheary element connection in the Arabidopsis/Nicotiana interfamilial grafts. Front. Plant Sci. 2021: 12: 664342.

Endo M, Mikami M, Endo A, Kaya H, Itoh T, Nishimasu H, Nureki O, Toki S. Genome editing in plants by engineered CRISPR-Cas9 recognizing NG PAM. Nat. Plants 2019: 5: 14–17.

Estavillo GM, Crisp PA, Pornsiriwong W, Wirtz M, Collinge D, Carrie C, Giraud E, Whelan J, David P, Javot H, et al. Evidence for a SAL1-PAP chloroplast retrograde pathway that functions in drought and high light signaling in Arabidopsis. Plant Cell 2011: 23: 3992–4012.

Fang Y, Qin X, Liao Q, Du R, Luo X, Zhou Q, Li Z, Chen H, Jin W, Yuan Y, et al. The genome of homosporous maidenhair fern sheds light on the euphyllophyte evolution and defences. Nat. Plants 2022: 8: 1024–1037.

Feng X, Liu W, Cao F, Wang Y, Zhang G, Chen ZH, Wu F. Overexpression of *HvAKT1* improves drought tolerance in barley by regulating root ion homeostasis and ROS and NO signaling. J. Exp. Bot. 2020: 71: 6587–6600.

Fouracre JP, Harrison CJ. How was apical growth regulated in the ancestral land plant? Insights from the development of non-seed plants. Plant Physiol. 2022: 190: 100–112.

Frangedakis E, Marron AO, Waller M, Neubauer A, Tse SW, Yue Y, Ruaud S, Waser L, Sakakibara K, Szovenyi P. What can hornworts teach us? Front. Plant Sci. 2023: 14: 1108027.

Fu L, Wu D, Zhang X, Xu Y, Kuang L, Cai S, Zhang G, Shen Q. Vacuolar H^+^-pyrophosphatase HVP10 enhances salt tolerance via promoting Na^+^ translocation into root vacuoles. Plant Physiol. 2022: 188: 1248–1263.

Geng Y, Yan A, Zhou Y. Positional cues and cell division dynamics drive meristem development and archegonium formation in Ceratopteris gametophytes. Commun. Biol. 2022: 5: 650.

Hassan MM, Zhang Y, Yuan G, De K, Chen J-G, Muchero W, Tuskan GA, Qi Y, Yang X. Construct design for CRISPR/Cas-based genome editing in plants. Trends Plant Sci. 2021: 26: 1133–1152.

He Y, Mudgett M, Zhao Y. Advances in gene editing without residual transgenes in plants. Plant Physiol. 2022: 188: 1757–1768.

Hetherington AM, Woodward FI. The role of stomata in sensing and driving environmental change. Nature 2003: 424: 901–908.

Huang X, Wang W, Gong T, Wickell D, Kuo LY, Zhang X, Wen J, Kim H, Lu F, Zhao H, et al. The flying spider-monkey tree fern genome provides insights into fern evolution and arborescence. Nat. Plants 2022: 8: 500–512.

Ishiga Y, Watanabe M, Ishiga T, Tohge T, Matsuura T, Ikeda Y, Hoefgen R, Fernie AR, Mysore KS. The SAL-PAP chloroplast retrograde pathway contributes to plant immunity by regulating glucosinolate pathway and phytohormone signaling. Mol Plant Microbe Interact 2017: 30: 829–841.

Ishizaki K, Chiyoda S, Yamato KT, Kohchi T. *Agrobacterium*-mediated transformation of the haploid liverwort *Marchantia polymorpha* L., an emerging model for plant biology. Plant Cell Physiol. 2008: 49: 1084–1091.

Ishizaki K, Nishihama R, Yamato KT, Kohchi T. Molecular genetic tools and techniques for *Marchantia polymorpha* research. Plant Cell Physiol. 2016: 57: 262–270.

Jia Q, Kong D, Li Q, Sun S, Song J, Zhu Y, Liang K, Ke Q, Lin W, Huang J. The function of inositol phosphatases in plant tolerance to abiotic stress. Int. J. Mol. Sci. 2019: 20: 3999.

Jiang W, He J, Babla M, Wu T, Tong T, Riaz A, Zeng F, Qin Y, Chen G, Deng F, et al. Molecular evolution and interaction of 14-3-3 proteins with H^+^-ATPases in plant abiotic stresses. J. Exp. Bot. 2024: 75: 689–707.

Jiang W, Pan R, Wu C, Xu L, Abdelaziz ME, Oelmüller R, Zhang W. *Piriformospora indica* enhances freezing tolerance and post-thaw recovery in Arabidopsis by stimulating the expression of *CBF* genes. Plant Signal Behav 2020: 15: 1745472.

Jiang W, Tong T, Li W, Huang Z, Chen G, Zeng F, Riaz A, Amoanimaa-Dede H, Pan R, Zhang W, et al. Molecular evolution of plant 14-3-3 proteins and function of Hv14-3-3A in stomatal regulation and drought tolerance. Plant Cell Physiol. 2022: 63: 1857–1872.

Kawai-Toyooka H, Kuramoto C, Orui K, Motoyama K, Kikuchi K, Kanegae T, Wada M. DNA interference: a simple and efficient gene-silencing system for high-throughput functional analysis in the fern *Adiantum*. Plant Cell Physiol. 2004: 45: 1648–1657.

Kim K-H, Song K, Park J-M, Kim J-Y, Lee B-M. RNA-Seq analysis of gene expression changes related to delay of flowering time under drought stress in tropical maize. Applied Sciences 2021: 11: 4273.

Kinosian SP, Wolf PG. The biology of *C. richardii* as a tool to understand plant evolution. Elife 2022: 11: e75019.

Kis A, Hamar E, Tholt G, Ban R, Havelda Z. Creating highly efficient resistance against wheat dwarf virus in barley by employing CRISPR/Cas9 system. Plant Biotechnol. J. 2019: 17: 1004–1006.

Klink VP, Wolniak SM. Centrin is necessary for the formation of the motile apparatus in spermatids of *Marsilea*. Mol. Biol. Cell 2001: 12: 761–776.

Kor SD, Chowdhury N, Keot AK, Yogendra K, Chikkaputtaiah C, Sudhakar Reddy P. RNA Pol III promoters-key players in precisely targeted plant genome editing. Front. Genet. 2022: 13: 989199.

Ku Y-S, Koo NS-C, Li FW-Y, Li M-W, Wang H, Tsai S-N, Sun F, Lim BL, Ko W-H, Lam H-M. GmSAL1 hydrolyzes inositol-1,4,5-trisphosphate and regulates stomatal closure in detached leaves and ion compartmentalization in plant cells. PLoS One 2013: 8: e78181.

Kuang L, Shen Q, Chen L, Ye L, Yan T, Chen ZH, Waugh R, Li Q, Huang L, Cai S, et al. The genome and gene editing system of sea barleygrass provide a novel platform for cereal domestication and stress tolerance studies. Plant Commun. 2022: 3: 100333.

Kumar V, Charde V, Prasad SB, Gandhi Y, Mishra SK, Rawat H, Thakur A, Shakya SK, Ansari T, Babu G, et al. Therapeutic potential of evergreen maiden hair fern *Adiantum venustum* D. Don: A comprehensive review. Food Chem. Adv. 2023: 3: 100439.

Li FW, Brouwer P, Carretero-Paulet L, Cheng S, de Vries J, Delaux PM, Eily A, Koppers N, Kuo LY, Li Z, et al. Fern genomes elucidate land plant evolution and cyanobacterial symbioses. Nat. Plants 2018: 4: 460–472.

Li W, Teng F, Li T, Zhou Q. Simultaneous generation and germline transmission of multiple gene mutations in rat using CRISPR-Cas systems. Nat. Biotechnol. 2013: 31: 684–686.

Liu H, Ding Y, Zhou Y, Jin W, Xie K, Chen L-L. CRISPR-P 2.0: An improved CRISPR-Cas9 tool for genome editing in plants. Mol. Plant 2017: 10: 530–532.

Liu J, Chen J, Zheng X, Wu F, Lin Q, Heng Y, Tian P, Cheng Z, Yu X, Zhou K, et al. GW5 acts in the brassinosteroid signalling pathway to regulate grain width and weight in rice. Nat. Plants 2017: 3: 17043.

Liu S, Essemine J, Liu Y, Liu C, Zhang F, Xu Z, Qu M. PIF4 promotes water use efficiency during fluctuating light and drought resistance in rice. bioRxiv 2023: 2023.2003. 2002.530909.

Liu XH, Fan Y, Mak M, Babla M, Holford P, Wang FF, Chen G, Scott G, Wang G, Shabala S, et al. QTLs for stomatal and photosynthetic traits related to salinity tolerance in barley. BMC Genomics 2017: 18: 1–13.

Lorenzo CD, Debray K, Herwegh D, Develtere W, Impens L, Schaumont D, Vandeputte W, Aesaert S, Coussens G, De Boe Y, et al. BREEDIT: a multiplex genome editing strategy to improve complex quantitative traits in maize. Plant Cell 2023: 35: 218–238.

Luo J, Li S, Xu J, Yan L, Ma Y, Xia L. Pyramiding favorable alleles in an elite wheat variety in one generation by CRISPR-Cas9-mediated multiplex gene editing. Mol. Plant 2021: 14: 847–850.

Ma C, Liu M, Li Q, Si J, Ren X, Song H. Efficient *BoPDS* gene editing in cabbage by the CRISPR/Cas9 system. Hortic. Plant J. 2019: 5: 164–169.

Ma J, Chen J, Wang M, Ren YL, Wang S, Lei CL, Cheng ZJ, Sodmergen. Disruption of OsSEC3A increases the content of salicylic acid and induces plant defense responses in rice. J. Exp. Bot. 2018: 69: 1051–1064.

Manmathan H, Shaner D, Snelling J, Tisserat N, Lapitan N. Virus-induced gene silencing of *Arabidopsis thaliana* gene homologues in wheat identifies genes conferring improved drought tolerance. J. Exp. Bot. 2013: 64: 1381–1392.

Marchant DB, Chen G, Cai S, Chen F, Schafran P, Jenkins J, Shu S, Plott C, Webber J, Lovell JT, et al. Dynamic genome evolution in a model fern. Nat. Plants 2022: 8: 1038–1051.

Marchant DB, Sessa EB, Wolf PG, Heo K, Barbazuk WB, Soltis PS, Soltis DE. The C-Fern (*Ceratopteris richardii*) genome: insights into plant genome evolution with the first partial homosporous fern genome assembly. Sci. Rep. 2019: 9: 18181.

Muthukumar B, Joyce BL, Elless MP, Stewart CN, Jr. Stable transformation of ferns using spores as targets: *Pteris vittata* and *Ceratopteris thalictroides*. Plant Physiol. 2013: 163: 648–658.

Nekrasov V, Wang C, Win J, Lanz C, Weigel D, Kamoun S. Rapid generation of a transgene-free powdery mildew resistant tomato by genome deletion. Sci. Rep. 2017: 7: 482.

O’Carrigan A, Hinde E, Lu N, Xu XQ, Duan HL, Huang GM, Mak M, Bellotti B, Chen ZH. Effects of light irradiance on stomatal regulation and growth of tomato. Environ. Exp. Bot. 2014: 98: 65–73.

Pacesa M, Pelea O, Jinek M. Past, present, and future of CRISPR genome editing technologies. Cell 2024: 187: 1076–1100.

Pan R, Buitrago S, Feng Z, Abou-Elwafa SF, Xu L, Li C, Zhang W. *HvbZIP21*, a novel transcription factor from wild barley confers drought tolerance by modulating ROS scavenging. Front. Plant Sci. 2022: 13: 878459.

Patel VK, Das A, Kumari R, Kajla S. Recent progress and challenges in CRISPR-Cas9 engineered algae and cyanobacteria. Algal Res. 2023: 71

Pennisi E. Fern proteins show promise against crop pests. Science 2023: 382: 868–869.

Petlewski AR, Li FW. Ferns: The final frond-tier in plant model systems. Am. Fern J. 2019: 109: 192–211.

Plackett AR, Conway SJ, Hewett Hazelton KD, Rabbinowitsch EH, Langdale JA, Di Stilio VS. *LEAFY* maintains apical stem cell activity during shoot development in the fern *Ceratopteris richardii*. Elife 2018: 7: e39625.

Plackett AR, Huang L, Sanders HL, Langdale JA. High-efficiency stable transformation of the model fern species *Ceratopteris richardii* via microparticle bombardment. Plant Physiol. 2014: 165: 3–14.

Plackett AR, Rabbinowitsch EH, Langdale JA. Protocol: genetic transformation of the fern *Ceratopteris richardii* through microparticle bombardment. Plant Methods 2015: 11: 37.

Plackett ARG, Emms DM, Kelly S, Hetherington AM, Langdale JA. Conditional stomatal closure in a fern shares molecular features with flowering plant active stomatal responses. Curr. Biol. 2021: 31: 4560–4570.

Pohthmi S, Sharma PB. A review of nutritional and ethno-medicinal properties of *Diplazium esculentum* (Retzius) Swart: A wild vegetable fern. Medicinal Plants-International Journal of Phytomedicines and Related Industries 2023: 15: 261–269.

Poovaiah C, Phillips L, Geddes B, Reeves C, Sorieul M, Thorlby G. Genome editing with CRISPR/Cas9 in *Pinus radiata* (D. Don). BMC Plant Biol. 2021: 21: 363.

Pornsiriwong W, Estavillo GM, Chan KX, Tee EE, Ganguly D, Crisp PA, Phua SY, Zhao C, Qiu J, Park J, et al. A chloroplast retrograde signal, 3’-phosphoadenosine 5’-phosphate, acts as a secondary messenger in abscisic acid signaling in stomatal closure and germination. eLife 2017: 6: e23361.

Proost S, Mutwil M. CoNekT: an open-source framework for comparative genomic and transcriptomic network analyses. Nucleic Acids Res. 2018: 46: W133–W140.

Qiu CW, Yue M, Wang QQ, Fu MM, Li C, Wang Y, Wu F. Barley HOMOCYSTEINE METHYLTRANSFERASE 2 confers drought tolerance by improving polyamine metabolism. Plant Physiol. 2023: 193: 389–409.

Rahmatpour N, Kuo LY, Kang J, Herman E, Lei L, Li M, Srinivasan S, Zipper R, Wolniak SM, Delwiche CF, et al. Analyses of *Marsilea vestita* genome and transcriptomes do not support widespread intron retention during spermatogenesis. New Phytol. 2023: 237: 1490–1494.

Ren Q, Sretenovic S, Liu S, Tang X, Huang L, He Y, Liu L, Guo Y, Zhong Z, Liu G, et al. PAM-less plant genome editing using a CRISPR-SpRY toolbox. Nat. Plants 2021: 7: 25-33.

Ren Q, Zhong Z, Wang Y, You Q, Li Q, Yuan M, He Y, Qi C, Tang X, Zheng X, et al. Bidirectional promoter-based CRISPR-Cas9 systems for plant genome editing. Front. Plant Sci. 2019: 10: 1173.

Rossel JB, Walter PB, Hendrickson L, Chow WS, Poole A, Mullineaux PM, Pogson BJ. A mutation affecting *ASCORBATE PEROXIDASE 2* gene expression reveals a link between responses to high light and drought tolerance. Plant, Cell & Environment 2005: 29: 269–281.

Rutherford G, Tanurdzic M, Hasebe M, Banks JA. A systemic gene silencing method suitable for high throughput, reverse genetic analyses of gene function in fern gametophytes. BMC Plant Biol. 2004: 4: 6.

Sánchez-León S, Gil-Humanes J, Ozuna CV, Giménez MJ, Sousa C, Voytas DF, Barro F. Low-gluten, nontransgenic wheat engineered with CRISPR/Cas9. Plant Biotechnol. J. 2018: 16: 902–910.

Shabala S, Chen G, Chen ZH, Pottosin I. The energy cost of the tonoplast futile sodium leak. New Phytol. 2020: 225: 1105–1110.

Shen Q, Zhang S, Ge C, Liu S, Chen J, Liu R, Ma H, Song M, Pang C. Genome-wide association study identifies GhSAL1 affects cold tolerance at the seedling emergence stage in upland cotton (*Gossypium hirsutum* L.). Theor. Appl. Genet. 2023: 136: 27.

Shin D-J, Min J-H, Van Nguyen T, Kim Y-M, Kim CS. Loss of *Arabidopsis Halotolerance 2-like* (*AHL*), a 3′-phosphoadenosine-5′-phosphate phosphatase, suppresses insensitive response of *Arabidopsis thaliana ring zinc finger 1* (*atrzf1*) mutant to abiotic stress. Plant Molecular Biology 2019: 99: 363–377.

Shirzadian-Khorramabad R, Moazzenzadeh T, Sajedi RH, Jing HC, Hille J, Dijkwel PP. A mutation in Arabidopsis SAL1 alters its in vitro activity against IP_3_ and delays developmental leaf senescence in association with lower ROS levels. Plant Mol Biol 2022: 108: 549–563.

Stout SC, Clark GB, Archer-Evans S, Roux SJ. Rapid and efficient suppression of gene expression in a single-cell model system, *Ceratopteris richardii*. Plant Physiol. 2003: 131: 1165–1168.

Tang X, Zheng X, Qi Y, Zhang D, Cheng Y, Tang A, Voytas DF, Zhang Y. A single transcript CRISPR-Cas9 system for efficient genome editing in plants. Mol. Plant 2016: 9: 1088–1091.

Tansley C, Houghton J, Rose AME, Witek B, Payet RD, Wu T, Miller JB. CIPK-B is essential for salt stress signalling in Marchantia polymorpha. New Phytol. 2023: 237: 2210–2223.

Taparia Y, Dolui AK, Boussiba S, Khozin-Goldberg I. Multiplexed genome editing via an RNA polymerase II promoter-driven sgRNA Array in the Diatom *Phaeodactylum tricornutum*: Insights into the role of StLDP. Front. Plant Sci. 2022: 12: 784780.

Tavernier E-K, Lockwood E, Perroud P-F, Nogué F, McDaniel SF. Establishing CRISPR-Cas9 in the sexually dimorphic moss, Ceratodon purpureus. bioRxiv 2023: 10.1101/2023.1110.1125.562971.

Tripathi L, Ntui VO, Tripathi JN. CRISPR/Cas9-based genome editing of banana for disease resistance. Curr. Opin. Plant Biol. 2020: 56: 118–126.

Van Bel M, Silvestri F, Weitz EM, Kreft L, Botzki A, Coppens F, Vandepoele K. PLAZA 5.0: extending the scope and power of comparative and functional genomics in plants. Nucleic Acids Res. 2022: 50: D1468–D1474.

Wang C, Li Y, Wang N, Yu Q, Li Y, Gao J, Zhou X, Ma N. An efficient CRISPR/Cas9 platform for targeted genome editing in rose (*Rosa hybrida*). J. Integr. Plant Biol. 2023: 65: 895–899.

Wang C, Wang Z, Cai Y, Zhu Z, Yu D, Hong L, Wang Y, Lv W, Zhao Q, Si L, et al. A higher-yield hybrid rice is achieved by assimilating a dominant heterotic gene in inbred parental lines. Plant Biotechnol. J. 2024:

Wang J, Wu H, Chen Y, Yin T. Efficient CRISPR/Cas9-mediated gene editing in an interspecific hybrid poplar with a highly heterozygous genome. Front. Plant Sci. 2020: 11: 996.

Wang NH, Zhou XY, Shi SH, Zhang S, Chen ZH, Ali MA, Ahmed IM, Wang Y, Wu F. An miR156-regulated nucleobase-ascorbate transporter 2 confers cadmium tolerance via enhanced anti-oxidative capacity in barley. J. Adv. Res. 2023: 44: 23–37.

Wang P, Zhang J, Sun L, Ma Y, Xu J, Liang S, Deng J, Tan J, Zhang Q, Tu L, et al. High efficient multisites genome editing in allotetraploid cotton (*Gossypium hirsutum*) using CRISPR/Cas9 system. Plant Biotechnol. J. 2018: 16: 137–150.

Wang W, Li Y-Z, Fan X-W, Chen Q, Zhong H. A photoperiod-responsive protein compendium and conceptual proteome roadmap outline in maize grown in growth chambers with controlled conditions. PLoS One 2017: 12: e0174003.

Wei JZ, Lum A, Schepers E, Liu L, Weston RT, McGinness BS, Heckert MJ, Xie W, Kassa A, Bruck D, et al. Novel insecticidal proteins from ferns resemble insecticidal proteins from *Bacillus thuringiensis*. Proc Natl Acad Sci U S A 2023: 120: e2306177120.

Wilson PB, Estavillo GM, Field KJ, Pornsiriwong W, Carroll AJ, Howell KA, Woo NS, Lake JA, Smith SM, Millar AH, et al. The nucleotidase/phosphatase SAL1 is a negative regulator of drought tolerance in Arabidopsis. Plant J. 2009: 58: 299–317.

Withers KA, Falls K, Youngstrom CE, Nguyen T, DeWald A, Yarvis RM, Simons GP, Flanagan R, Bui LT, Irish EE, et al. A Ceratopteris EXCESS MICROSPOROCYTES1 suppresses reproductive transition in the fern vegetative leaves. Plant Sci. 2023: 335: 111812.

Xi H, Xu H, Xu W, He Z, Xu W, Ma M. A SAL1 loss-of-function Arabidopsis mutant exhibits enhanced cadmium tolerance in association with alleviation of endoplasmic reticulum stress. Plant Cell Physiol. 2016: 57: 1210–1219.

Xie K, Minkenberg B, Yang Y. Boosting CRISPR/Cas9 multiplex editing capability with the endogenous tRNA-processing system. Proc Natl Acad Sci U S A 2015: 112: 3570–3575.

Xiong L, Lee B-h, Ishitani M, Lee H, Zhang C, Zhu J-K. FIERY1 encoding an inositol polyphosphate 1-phosphatase is a negative regulator of abscisic acid and stress signaling in Arabidopsis. Genes & Development 2001: 15: 1971–1984.

Yan H, Gao Y, Wu L, Wang L, Zhang T, Dai C, Xu W, Feng L, Ma M, Zhu YG, et al. Potential use of the *Pteris vittata* arsenic hyperaccumulation-regulation network for phytoremediation. J. Hazard. Mater. 2019: 368: 386–396.

Ye S, Ding W, Bai W, Lu J, Zhou L, Ma X, Zhu Q. Application of a novel strong promoter from Chinese fir (*Cunninghamia lanceolate*) in the CRISPR/Cas mediated genome editing of its protoplasts and transgenesis of rice and poplar. Front. Plant Sci. 2023: 14: 1179394.

Youngstrom CE, Geadelmann LF, Irish EE, Cheng CL. A fern *WUSCHEL-RELATED HOMEOBOX* gene functions in both gametophyte and sporophyte generations. BMC Plant Biol. 2019: 19: 416.

Youngstrom CE, Withers KA, Irish EE, Cheng CL. Vascular function of the T3/modern clade WUSCHEL-Related HOMEOBOX transcription factor genes predate apical meristem-maintenance function. BMC Plant Biol. 2022: 22: 210.

Yu J, Lee KM, Son M, Kim KH. Effects of the deletion and over-expression of *Fusarium graminearum* gene *FgHal2* on host response to mycovirus *Fusarium graminearum virus 1*. Mol. Plant Pathol 2015: 16: 641–652.

Yu X, Meng X, Liu Y, Wang X, Wang T-J, Zhang A, Li N, Qi X, Liu B, Xu Z-Y. The chromatin remodeler ZmCHB101 impacts alternative splicing contexts in response to osmotic stress. Plant Cell Rep. 2018: 38: 131–145.

Yuan J, Xu T, Hiltbrunner A. Phytochrome higher order mutants reveal a complex set of light responses in the moss *Physcomitrium patens*. New Phytol. 2023: 239: 1035–1050.

Zhang J, Wu Q, Eleouet M, Chen R, Chen H, Zhang N, Hu Y, Sui Z. CRISPR/LbCas12a-mediated targeted mutation of *Gracilariopsis lemaneiformis* (Rhodophyta). Plant Biotechnol. J. 2023: 21: 235–237.

Zhao C, Wang Y, Chan KX, Marchant DB, Franks PJ, Randall D, Tee EE, Chen G, Ramesh S, Phua SY, et al. Evolution of chloroplast retrograde signaling facilitates green plant adaptation to land. Proc. Natl. Acad. Sci. U.S.A. 2019: 116: 5015–5020.

